# An epitope specific patient-derived LGI1-autoantibody enhances neuronal excitability by modulating the Kv1.1 channel

**DOI:** 10.1101/2021.12.10.471894

**Authors:** Johanna Extrémet, Oussama El Far, Sarosh R Irani, Dominique Debanne, Michael Russier

**Author notes:** Correspondence to Michael Russier.

## Abstract

Leucine-rich Glioma Inactivated protein 1 (LGI1) is expressed in the central nervous and genetic loss of function is associated with epileptic disorders. Also, patients with LGI1-directed autoantibodies have frequent focal seizures as a key feature of their disease. LGI1 is composed of a Leucine Rich Repeat (LRR) and an Epitempin (EPTP) domain. These domains are reported to interact with different aspects of the transsynaptic complex formed by LGI1 at excitatory synapses, including presynaptic Kv1 potassium channels. Patient-derived monoclonal antibodies (mAbs) are ideal reagents to study whether domain-specific LGI1-autoantibodies induce epileptiform activities in neurons, and their downstream mechanisms. To address this question, we measured the intrinsic excitability of CA3 pyramidal neurons in organotypic cultures from rat hippocampus treated with either a LRR- or an EPTP-reactive patient-derived mAb. The antibodies induced changes in neuronal intrinsic excitability which led us to measure their effects on Kv1-type potassium currents. We found an increase of intrinsic excitability correlated with a reduction of the sensitivity to a selective Kv1.1-channel blocker in neurons treated with the LRR mAb compared to the control, but not in neurons treated with the EPTP mAb. Our findings suggest LRR mAbs are able to modulate neuronal excitability that could account for epileptiform activities observed in patients.

## Introduction

Autoantibodies directed against Leucine-rich Glioma-Inactivated protein 1 (LGI1) are found in patients with limbic encephalitis (LE) who have frequent focal seizures and hippocampal atrophy as hallmarks of their disease (Aurangzeb et al., 2017; Irani et al., 2011; Thompson et al., 2018). Hyperexcitability and epileptiform activities have been recorded in the hippocampi of mice and rats treated with serum from patients with LGI1-autoantibodies, suggesting their direct pathogenicity (Lalic et al., 2011; Petit-Pedrol et al., 2018). LGI1 is a soluble molecule composed of a Leucine-Rich Repeat (LRR) and an Epitempin (EPTP) domain. The LRR domain is reported to mediate transsynaptic homo-oligomerisation whereas the EPTP domain allows LGI1 to dock on its pre- and post-synaptic receptors, ADAM23 and ADAM22, respectively (Yamagata and Fukai, 2020). Serum LGI1-antibodies have been shown to target both the LRR and EPTP domains of LGI1. Sera have been shown to cause both a down-regulation of two interaction partners of LGI1: the presynaptic voltage-gated potassium channel Kv1 and the postsynaptic AMPA receptors (Kornau et al., 2020; Lalic et al., 2011; Petit-Pedrol et al., 2018; Ramberger et al., 2020). Furthermore, polyclonal serum antibodies can prevent the interaction of LGI1 with ADAM22 and potentiate excitatory glutamatergic synapses (Kornau et al., 2020; Ohkawa et al., 2013; Ramberger et al., 2020).

More recent reports aimed to better characterize the relative pathogenic roles of domain-specific antibodies by studying monoclonal LGI1-autoantibodies (mAb) derived from patient B cells (Kornau et al., 2020; Ramberger et al., 2020). mAbs targeting the LRR domain were shown to internalise the LGI1 protein after it docked to ADAM22/ADAM23 receptors (Ramberger et al., 2020). In contrast, EPTP-mAbs operated by competing with LGI1 for binding to ADAM22 (Kornau et al., 2020; Ramberger et al., 2020). Both mAb categories were shown to modulate neuronal excitability, but LRR-mAbs induced more robust effects and behavioural deficits (Kornau et al., 2020; Ramberger et al., 2020).

In order to directly compare the electrophysiolgical effects of LRR- and EPTP-mAbs, we measured intrinsic excitability of CA3 neurons after their application. As LGI1 can alter the inactivation gating of the Kv1 channels (Schulte et al., 2006) which tune neuronal excitability in CA3 pyramidal neurons (Rama et al., 2017), we tested whether the blocking the Kv1.1 subunit with a specific antagonist, dendrotoxin-k (DTx-k), affected intrinsic excitability which could be abrogated by the mAbs.

## Materials & Methods

### Rat hippocampal slice cultures

All experiments were carried out according to the European and Institutional guidelines for the care and use of laboratory animals and approved by the local health authority (D13055-08, Préfecture des Bouches-du-Rhône). Slice cultures were prepared as described previously (Debanne et al., 2008). In brief, young Wistar rats (P7–P10) were anesthetized with isoflurane and killed by decapitation, the brain was removed, and each hippocampus was dissected. Hippocampal slices (350 μm) were obtained using a Vibratome (Leica, VT1200S). They were placed on 20-mm latex membranes (Millicell) inserted into 35-mm Petri dishes containing 1 mL of culture medium and maintained for up to 21 d in an incubator at 34°C, 95% O_2_–5% CO_2_. The culture medium contained 25ml MEM, 12.5ml HBSS, 12.5ml horse serum, 0.5ml penicillin/streptomycin, 0.8ml glucose (1 M), 0.1ml ascorbic acid (1 mg/mL), 0.4ml Hepes (1 M), 0.5ml B27, and 8.95ml sterile H_2_O. To limit glial proliferation, 5 μM Ara-C was added to the culture medium starting at 4 d in vitro (DIV).

### Application of the mAbs

LGI1-specific mAbs were expressed in HEK293T cells, as described in Ramberger et al. 2020. Polyclonal IgGs purified from total healthy human serum (Protein G Sepharose 4 Fast Flow, 17-0618-01, Cytiva) were used as a control (Human Control: HC). The electrophysiological effect of one recombinant LRR-mAb (mAb01, Ramberger et al., 2020) and one EPTP-mAb (mAb6212) were assessed individually in rat CA3 pyramidal neurons. HC IgGs and LGI1-mAbs were applied daily from DIV4 until the day of the experiment, at a final concentration of 4 ng/μl in culture medium, in direct contact with the hippocampal organotypic cultures.

### Electrophysiology

Whole-cell recordings were obtained from CA3 neurons in organotypic cultures at DIV7 and DIV8. Recordings were carried out in a limit of 6 h after the last application of the IgGs. The external solution contained: 125mM NaCl, 26mM NaHCO_3_, 3mM CaCl_2_, 2.5mM KCl, 2mM MgCl_2_, 0.8mM NaH_2_PO_4_, and 10mM D-glucose, and was equilibrated with 95% O_2_–5% CO_2_. Patch pipettes (7– 10 MΩ) were filled with a solution containing (mM): K-gluconate 120, KCl 20, Hepes 10, EGTA 0.5, MgCl_2_ 2, Na_2_ATP 2, and NaGTP 0.3 (pH 7.4). Organotypic cultures were quickly washed in the perfused recording chamber with the external solution by increasing the speed of perfusion pumps to facilitate the patch-clamp procedure. All recordings were made at 29°C in a temperature-controlled recording chamber (Luigs & Neumann). Neurons were recorded in current clamp with a Multiclamp 700B Amplifier (Axon Instruments, Molecular Devices). Excitability was measured by delivering a range of long (1 s) depolarizing current pulses (10–250 pA, by increments of 10 pA) and counting the number of action potentials. Ionotropic glutamate and GABA receptors were blocked with 2–4 mM kynurenate and 100 μM picrotoxin, respectively. Input–output curves corresponding to the number of action potential elicited by each increment of injected current were determined for each neuron and two parameters were examined; the rheobase (the minimal current eliciting at least one action potential) and the first spike latency (depolarising time before the first spike obtained under rheobase current).

Sensitivity of intrinsic excitability to the selective Kv1.1-channel blocker, dendrotoxin-K (DTx-k) was determined by current-clamp recording before and after 5 minutes of bath application of DTx-K (100 nM). Acquisition was performed at 10 kHz with pClamp10 (Axon Instruments).

Data were analysed with ClampFit (Axon Instruments) and IgorPro (Wavemetrics). Pooled data are presented as mean ± SEM and statistical analysis was performed using the Mann–Whitney U test or Wilcoxon rank signed test.

## Results

### The LRR-mAb, but not the EPTP-mAb, increases intrinsic excitability in CA3 pyramidal neurons

To assess the effect of the two mAbs on neuronal excitability, we performed current-clamp recordings from CA3 pyramidal neurons in treated organotypic cultures of rat hippocampus. A significant decrease in the rheobase was observed after treatment with the LRR-mAb (HC IgGs: 100.1 ± 7.8 pA n = 13 vs. LRR-mAb: 75.4 ± 7.5 pA n = 14, Mann-Whitney, p = 0.035) but not with the EPTP-mAb (HC IgGs: 100.1 ± 7.8 pA n = 13 vs. EPTP-mAb: 110.2 ± 15.6 pA n = 12, Mann-Whitney, p = 0.96) (Figure 1A). In addition, CA3 neurons treated with the LRR-mAb showed a leftward shift of the input-output curve compared to neurons treated with HC IgGs, whereas input-output curves were not different between neurons treated with the EPTP-mAb and HC IgGs (Figure 1B). These data suggest LRR-mAbs could induce an increase in intrinsic excitability in rat CA3 neurons without differences in passive properties of recorded neurons (input resistance: 225.85 ± 7.37 MΩ n = 13 vs. LRR-mAb: 222.45 ± 13.09 MΩ, n = 14; Mann-Whitney, p = 0.45 or capacitance: 337.85 ± 14.11 pF n = 13 vs. LRR-mAb: 329.92 ± 20.45 pF n = 14, Mann-Whitney, p = 0.68, data not shown).

**Fig. 1.**
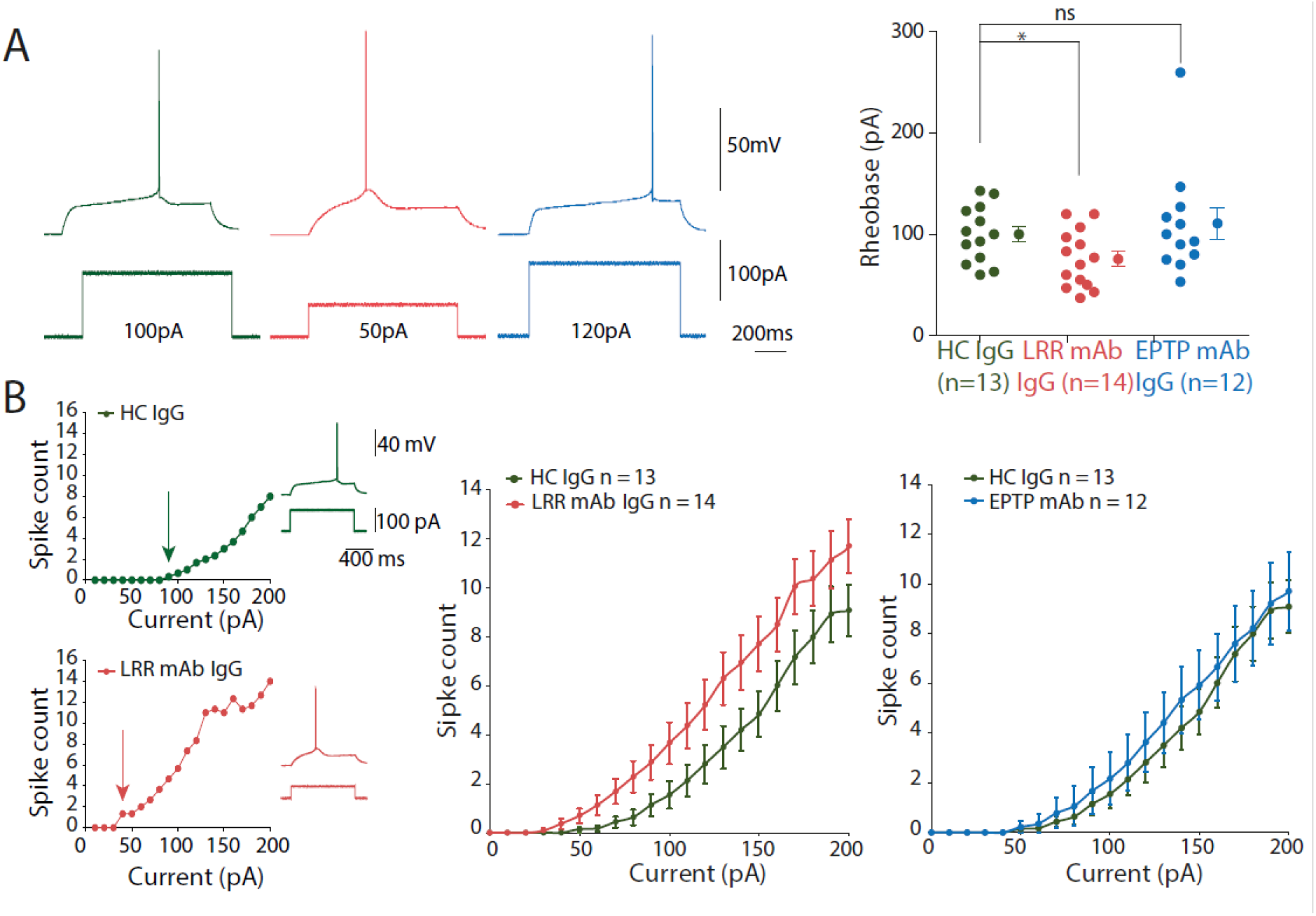
Application of patient-derived LRR-mAb increases intrinsic excitability of hippocampal CA3 neurons. (A, Left) Traces illustrate the rheobase current of neurons from rat hippocampal organotypic cultures treated with the HC IgGs (green), LRR-mAb (red) and EPTP-mAb (blue). (A, Right) Graph of rheobase current is represented with the mean value for each condition. (B, Left) Input-output curves showing the number of spikes according to depolarising current increments and corresponding traces of representative neurons treated with the HC IgG (Top) and LRR-mAb (Bottom) recorded in current clamp. The rheobase is indicated by arrows and illustrated by the traces. (B, Right) Averaged input-output curves of the two mAbs are compared to control IgGs. Error bars represent SEM. Statistical analysis was performed using the Mann-Whitney test (graph A) and significance was obtained at P<0.05.

### The LRR-mAb increases intrinsic excitability through the modulation of the Kv1 potassium current

Recordings of HC IgGs-treated neurons showed a characteristic ramp-and-delay profile preceding the evoked spike, strikingly observed at rheobase current (Figure 2A). This feature also found in untreated control neurons was characterized as the hallmark of fast-activating slow offset conductance of the D-type current carried by the voltage-gated potassium Kv1 channel (Cudmore et al., 2010; Seagar et al., 2017). Interestingly, this profile was replaced by a rapid depolarization eliciting the first spike in CA3 neurons treated with the LRR-mAb (Figure 2A). To evaluate this difference, we measured the latency to the first spike. CA3 neurons treated with the LRR-mAb showed a reduced first spike latency compared to HC IgGs treated neurons (494 ± 42 ms n = 14 vs 728 ± 43 ms n = 13, Mann-Whitney, p = 0.001; Figure 2A) without significant difference between neurons treated with HC IgGs or EPTP-mAb (728 ± 43 ms n = 13 vs. 762 ± 57 ms n = 12, Mann-Whitney, ns; Figure 2A).

**Fig. 2.**
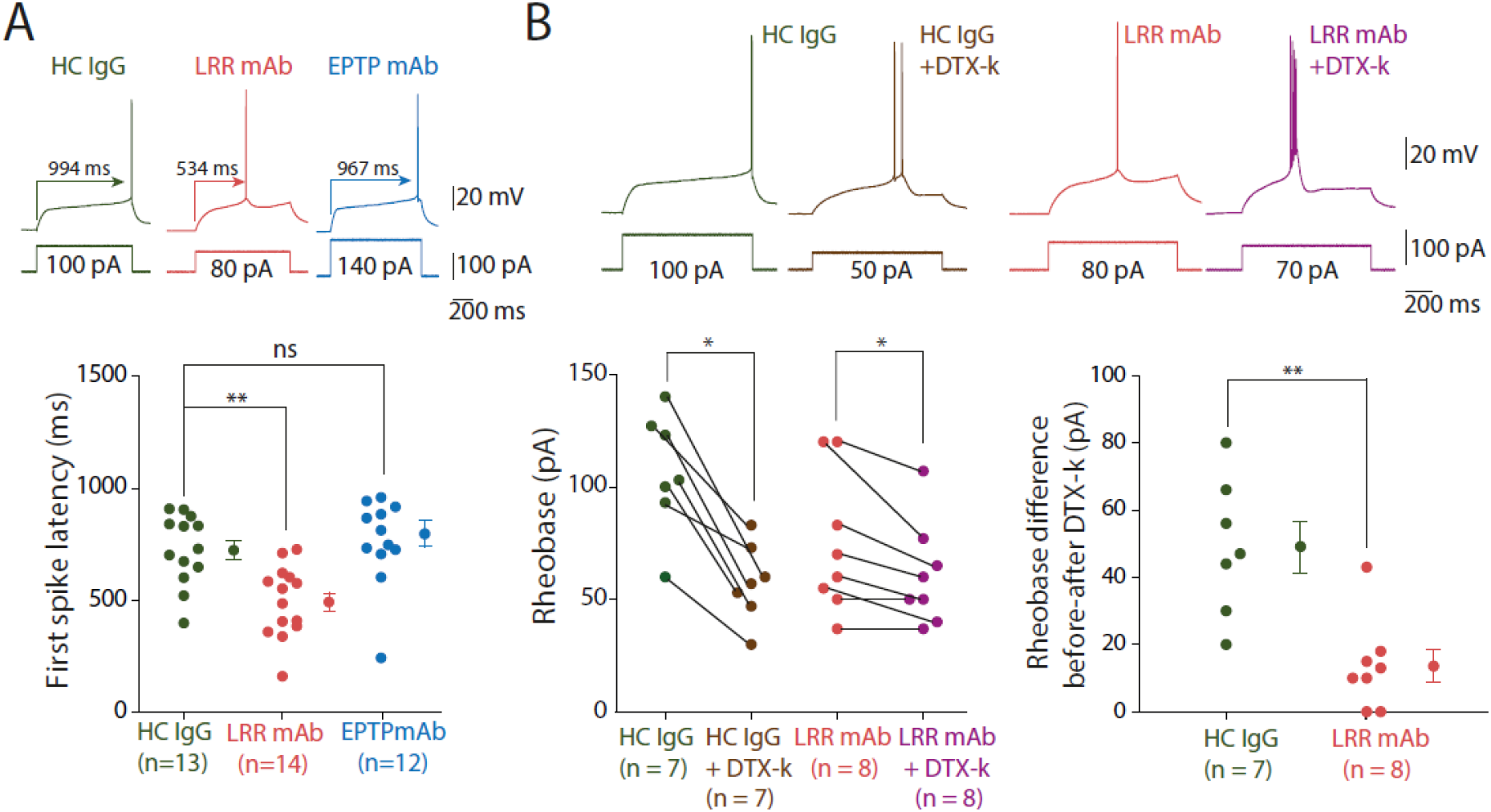
LRR-mAb-mediated increase of intrinsic excitability in hippocampal CA3 neurons is induced by a decrease of D-type current. (A) The latency to the first spike is illustrated by the traces at the rheobase current and values are reported in the graph for neurons treated with HC IgGs (green), LRR-mAb (red) and EPTP-mAb (blue). (B, Left) Graph and traces shows the rheobase current before and after bath application of DTX-k (100 nM) after HC IgGs treatment (green and brown) and after LRR-mAb treatment (red and purple). (B, Right) Graph highlighting the effect of DTX-k by the subtraction of the rheobase after adding DTX-k to the rheobase before DTX-k, after each antibody treatment. Error bars represent SEM. Statistical analysis was performed using the Mann-Whitney test (A and B Right) and the Wilcoxon rank-signed test (B Left) and significance was obtained at P<0.05.

Based on these findings, we asked whether changes in the D-type potassium current could account for the change in intrinsic excitability by measuring the sensitivity to DTx-k after LRR-mAb treatment. In contrast to HC IgG treated neurons (rheobase = 107 ± 10 pA vs. 58 ± 6 pA n = 7, Wilcoxon, p = 0.016; Figure 2B), or neurons treated with the EPTP-mAb (rheobase = 115 ± 23 pA vs. 63 ± 12 pA n = 9, Wilcoxon, p = 0.009; data not shown) which displayed a robust decrease in rheobase after bath application of DTx-k, a modestly significant effect of DTx-k was observed in neurons treated with the LRR-mAb (74 ± 11 pA vs. 61 ± 8 pA n = 8, Wilcoxon, p = 0.035; Figure 2B). Moreover, bath application of DTx-k induced a loss of the ramp-and-delay phenotype only in neurons treated with HC IgGs whereas no difference was observed in neurons treated with the LRR-mAb (Figure 2B). Taken together, these data suggest that the increase of intrinsic excitability measured in CA3 neurons treated with the LRR-mAb may be mediated by a reduction of D-type current. To confirm this, we calculated the effect of LRR-mAb on sensitivity to DTx-k with the rheobase difference before and after adding DTx-k. Whereas no significant difference was observed between neurons treated with HC IgGs or EPTP-mAb (49 ± 8 pA n = 7 vs. 52 ± 16 pA n = 9 respectively, Wilcoxon, p = 0.56; data not shown), we found a 5 fold decrease in rheobase difference in neurons treated with the LRR-mAb compared to HC IgGs (14 ± 5 pA n = 8 vs. 49 ± 8 pA n = 7, respectively Mann-Whitney, p = 0.003; Figure 2B). These data confirm that a loss of a D-type current induced by the LRR-mAb contributes significantly to the observed increase in intrinsic excitability.

## Discussion

In our study, we investigated whether two patients–derived mAbs directed against different epitopes on LGI1 perturb neuronal excitability. From rat hippocampal organotypic cultures, input-output curves and rheobase currents showed an increase in intrinsic excitability after application of the LRR-mAb but not with the EPTP-mAb. Further, treatment with the LRR-mAb, but not the EPTP-mAb, prevented the effect of DTX-k, a blocker of the Kv1.1 voltage-gated potassium channel. LRR-mAbs have been shown to promote LGI1-ADAM complex internalisation (Ramberger et al., 2020) which may lead to a reduction in Kv1 channel expression at the cell membrane. This mechanism is supported by genetic deletion of LGI1 which decreases Kv1.1 density and D-type currents by more than 50% (Seagar et al., 2017). This decrease in D-type currents is likely to increase neuronal excitability.

The molecular mechanism by which EPTP-mAbs mediate their effect is likely to occur through the prevention of LGI1 binding to its native receptors. Very weak binding of anti-EPTP mAbs to human LGI1 expressed with ADAM22/ADAM23 in HEK293T and rat hippocampal cultures or on mouse brain sections has been reported (Kornau et al., 2020; Ramberger et al., 2020). Even if differences in peptide sequences between rat LGI1 and its human homologue could determine the ability of the antibody to have a functional effect, a lack of epitope availability by a competition between EPTP antibodies and ADAM22/ADAM23 in these systems could be an hypothesis to explore (Ohkawa et al., 2013; Ramberger et al., 2020). In our study, EPTP epitope of endogenous LGI1 could be occupied by its receptors which could prevent EPTP-mAb to have a functional effect. Further experiments are needed to elucidate these questions.

## Acknowledgments

This work was supported by INSERM, CNRS, ANR (ANR-17-CE16-0022 LoGiK to DD and OEF).

## Author contribution

OEF, MR and DD designed research; JE performed the experiments; SI provided the antibodies; JE, DD & MR analysed the data; JE, DD, OEF & MR wrote the paper, which was approved by all the authors.

